# Parsing heterogeneity in autism spectrum disorder and attention-deficit/hyperactivity disorder with individual connectome mapping

**DOI:** 10.1101/490672

**Authors:** Dina R. Dajani, Catherine A. Burrows, Mary Beth Nebel, Stewart H. Mostofsky, Kathleen M. Gates, Lucina Q. Uddin

## Abstract

Traditional diagnostic systems for mental illnesses define diagnostic categories that are heterogeneous in behavior and underlying neurobiological alterations. The goal of this study was to parse heterogeneity in a core executive function, cognitive flexibility, in children with a range of abilities (*N*=132; children with autism spectrum disorder [ASD], attention deficit/hyperactivity disorder [ADHD], and typically developing [TD] children) using directed functional connectivity profiles derived from resting-state fMRI data. Brain regions activated in response to a cognitive flexibility task in adults were used guide region-of-interest (ROI) selection to estimate individual connectivity profiles in this study. We expected to find at least three subgroups of children who differed in their network connectivity metrics and symptom measures. Unexpectedly, we did not find a stable or valid subgrouping solution, which suggests that categorical models of the neural substrates of cognitive flexibility in children may be invalid. Results shed light on the validity of conceptualizing the neural substrates of cognitive flexibility categorically in children. Ultimately, this work may provide a foundation for the development of a revised nosology focused on neurobiological substrates of mental illness as an alternative to traditional symptom-based classification systems.

---

Autism spectrum disorder (ASD) and attention-deficit/hyperactivity disorder (ADHD) are prevalent neurodevelopmental disorders most commonly diagnosed in the United States according to a symptom-based classification system, the Diagnostic and Statistical Manual of Mental Health Disorders 5 (DSM-5, American Psychiatric Association 2013). Although these disorders are characterized by separate core deficits (in ASD, social communication deficits and restricted and repetitive behaviors [RRBs]; in ADHD, primarily inattentive and/or hyperactive and impulsive symptoms), overlap in behavioral presentation and biological substrates obfuscate the distinctions between these diagnostic categories. In particular, there is significant variability among children with ASD and ADHD concerning deficits in a subdomain of executive function (EF) – cognitive flexibility (Dajani, Llabre et al. 2016). Mixed evidence for distinct diagnostic categories suggests an alternative diagnostic system focusing on the full range of variation in behavior (i.e., Research Domain Criteria [RDoC], Cuthbert and Insel 2013) may be better-suited to identify individuals with a common biological pathway to abnormality. As a first step towards developing an improved nosology for neurodevelopmental disorders, this study aims to use individual connectome mapping to identify children with altered brain network connectivity that may contribute to impaired cognitive flexibility.

Cognitive flexibility undergoes protracted development across childhood to young adulthood (Anderson 2008), supporting a wide range of behaviors that impact life outcomes (Diamond and Lee 2011). In childhood, effective cognitive flexibility predicts better math and reading abilities (Bull, Espy et al. 2008) and social-emotional development such as false belief understanding (Farrant, Maybery et al. 2012). Deficits in cognitive flexibility can occur in healthy children and adults without accompanying mental illness, but deficits are more prevalent in almost every psychiatric population, such as autism spectrum disorder, attention-deficit/hyperactivity disorder, conduct disorder, depression, obsessive-compulsive disorder, substance abuse and schizophrenia (Diamond and Lee 2011, McTeague, Goodkind et al. 2016). Heightened rigidity in cognition and behavior may predispose individuals to poorer psychiatric morbidity, as evidenced by higher relapse rates in schizophrenia (Chen, Hui et al. 2005). On the other hand, behavioral interventions that improve cognitive flexibility have shown to also ease associated psychiatric symptoms (Tamm, Nakonezny et al. 2014). The ramifications of deficits in cognitive flexibility cannot be ignored. It is therefore imperative that we understand the neural underpinnings of this essential skill across the lifespan.

Traditional diagnostic systems for mental illnesses are limited in that they define diagnostic categories that have high biological and behavioral heterogeneity, allow for high between-category overlap, and cannot accurately predict treatment responsiveness or prognosis. Person-centered studies, which take into account within-population heterogeneity, substantiate evidence that the majority of (but not all) children with ASD and ADHD exhibit severe cognitive flexibility deficits (Gioia, Isquith et al. 2002, Fair, Bathula et al. 2012). Deficits in cognitive flexibility are particularly concerning because they are related to higher levels of core symptomatology in both ASD and ADHD: elevated RRBs in children with ASD and worse hyperactive-impulsive symptoms, oppositional defiant disorder symptoms, and lower intelligence in children with ADHD (Lopez, Lincoln et al. 2005, D’Cruz, Ragozzino et al. 2013, Roberts, Martel et al. 2013). In addition, there are high rates of comorbidity between diagnostic groups, with rates of comorbid ADHD in children with ASD ranging from 37-85% (Leitner 2014). Numerous studies have demonstrated the detrimental impact of comorbidity between ASD and ADHD diagnoses, citing poorer adaptive functioning, health-related quality of life (Sikora, Vora et al. 2012) and higher rates of clinically impaired cognitive inflexibility than children with either disorder alone (Yerys, Wallace et al. 2009, Dajani, Llabre et al. 2016). Further, there is strong evidence for shared heritability of ASD and ADHD (Ghirardi, Brikell et al. 2018). An alternative nosology based on neurobiologically homogeneous subgroups will aid in the ultimate goal to identify children who stand to benefit from targeted treatments specific to their brain network connectivity alterations (Cuthbert 2014). Due to the controversies surrounding the separability of ASD and ADHD diagnostic categories based on both biological and behavioral characteristics, we propose that children with one disorder or the other should be considered together in an effort to develop an alternative nosology. This conceptualization is in line with the RDoC approach, which advocates for single studies to span multiple diagnostic groups (Cuthbert 2014).

One unresolved issue in psychiatry is whether psychopathology should be conceptualized categorically, similar to traditional systems, or dimensionally, including the full range of behavior from normal to abnormal (Coghill and Sonuga-Barke 2012). While a categorical approach implies that mental disorders are qualitatively different from typical behavior, dimensional approaches assume that mental disorders are an extreme on a continuum of behavior represented across the population. Although many studies have attempted to find homogeneous, categorical subgroups within ASD and ADHD diagnostic categories using psychological or brain imaging data (e.g., van der Meer, Oerlemans et al. 2012, Gates, Molenaar et al. 2014, Costa Dias, Iyer et al. 2015, Kernbach, Satterthwaite et al. 2018), very few studies explicitly compare categorical and dimensional fits of the data using taxometric or factor mixture modeling. These few available studies have only focused on symptom-based and neuropsychological measures, demonstrating that ASD is a discrete category distinct from typical social communication and repetitive behaviors (Frazier, Youngstrom et al. 2010, Frazier, Youngstrom et al. 2012) and suggest the presence of three subgroups within the ASD category (Georgiades, Szatmari et al. 2013). Conversely, studies have consistently demonstrated a dimensional structure for ADHD symptoms in children (Haslam, Williams et al. 2006, Frazier, Youngstrom et al. 2007, Lubke, Hudziak et al. 2009, Marcus and Barry 2011), though one notable exception found a categorical fit using measures of EF and reward processing tasks (Stevens, Pearlson et al. 2018). Other studies focusing on resting state functional connectivity of large-scale brain networks report that ASD and ADHD symptomatology comprise both categorical and dimensional aspects, but did not explicitly test model fit (Chabernaud, Mennes et al. 2012, Elton, Alcauter et al. 2014, Elton, Di Martino et al. 2016). Therefore, it remains an open question whether the structure of neurodevelopmental disorders is categorical or dimensional at the neurobiological level.

One well-replicated finding in clinical psychology is a hierarchical taxonomy of psychopathology that exists dimensionally across healthy and patient populations (Lahey, Krueger et al. 2017). This ‘p’ factor encapsulates the propensity to develop any form of psychopathology, including anxiety, depression, substance use disorders, and schizophrenia. In individuals with a diagnosed mental health disorder, the ‘p’ factor describes the severity, chronicity, and comorbidity of symptoms. In children, the ‘p’ factor extends to symptoms characteristic of neurodevelopmental disorders, including ADHD and ASD (Neumann, Pappa et al. 2016, Martel, Pan et al. 2017). Further, in children, this general liability to develop psychopathology has been related to poorer EF abilities, poorer effortful control, and is heritable via single-nucleotide polymorphisms (Neumann, Pappa et al. 2016, Martel, Pan et al. 2017). Dimensional structures at the symptom-level do not necessarily imply the same structure at the brain network-level, but initial neuroimaging findings support dimensional models of psychopathology. Dysfunctional activation and functional connectivity of networks important for EF underlie this ‘p’ factor in children and adolescents (Shanmugan, Wolf et al. 2016, Liu, Liao et al. 2018).

Similar to the ‘p’ factor models, the RDoC framework opts to model behavior dimensionally, but moves away from focusing on DSM symptoms and towards neurobiologically validated functional constructs, such as EF (Garvey, Avenevoli et al. 2016). Deficits in EF are a transdiagnostic marker of psychopathology, meaning deficits can be detected in children without any traditional diagnoses, and across diagnostic groups such as ASD and ADHD (Dajani, Llabre et al. 2016, McTeague, Goodkind et al. 2016). The current study focuses on a specific aspect of EF, cognitive flexibility, because cognitive flexibility is impacted in various neurodevelopmental disorders (Dajani, Llabre et al. 2016) and these impairments relate to worse symptom severity and lower academic achievement in youth (Lopez, Lincoln et al. 2005, D’Cruz, Ragozzino et al. 2013, Roberts, Martel et al. 2013). Moreover, established flexibility-specific interventions exist that improve deficits in children with ASD and ADHD (Kenworthy, Anthony et al. 2014, Tamm, Nakonezny et al. 2014). EF deficits that are left untreated may persist and impact social competence and friendship quality in adolescence (Rosenthal, Wallace et al. 2013, Lieb and Bohnert 2017). Thus, it is imperative that researchers develop a reliable method to identify children who exhibit the most severe flexibility deficits.

As a basis for an alternative nosology, we leverage the strength of neurobiological variables due to the many limitations of behavior-based classifications (Waterhouse and Gillberg 2014). Most notably, behavior does not map one-to-one to underlying neurobiology (Pessoa 2014), meaning many disparate brain alterations may manifest as a singular phenotype (referred to as redundancy, see Licinio and Wong 2013). This is especially problematic if treatments only benefit subgroups of children with similar underlying neurobiological deficits (Loth, Spooren et al. 2016). Our group recently demonstrated that subgrouping approaches applied only to behavioral variables do not map well onto neurobiologically defined metrics. We developed an EF-based classification system based on parent-report and behavioral measures. This behavior-based classification system did *not* produce neurobiologically distinct subgroups, as assessed by functional connectivity metrics indexing the integrity of large-scale functional brain networks important for cognition (Dajani, Burrows et al., https://www.biorxiv.org/content/early/2018/08/22/396317). These data underscore the importance of moving beyond behavioral observations and focusing to a greater extent on differences in underlying *neurobiological* mechanisms of these disorders. Therefore, in this study we leverage large-scale neural networks as the foundation for developing an alternative nosology for two prevalent neurodevelopmental disorders.

Here, we focus on functional connectivity approaches estimated with resting-state fMRI data due to the large body of work implicating fronto-parietal functional networks in cognitive flexibility (Kim, Cilles et al. 2012). Specifically, we aim to investigate functional brain networks important for cognitive flexibility using the triple network model of psychopathology (Menon 2011), focusing on the frontoparietal (FPN), salience (SN), and default mode networks (DMN). We use a sample of children with ASD, ADHD and TD children with a wide range of EF abilities to identify subgroups with similar brain network connectivity profiles or “connectomes”. This study applies a cutting-edge method to construct individual-level connectomes by combining structural equation modeling (SEM) and unsupervised machine learning to rs-fMRI data called Group Iterative Multiple Model Estimation (GIMME, Gates and Molenaar 2012).

Most studies intending to characterize the neurobiological underpinnings of ASD and ADHD do not aim to understand heterogeneity inherent to each disorder, but instead effectively treat this variability as noise (Lenroot and Yeung 2013). As a result, only two published studies have investigated subgroups of children with ADHD based on functional connectivity metrics, and no study has examined brain network connectivity-based subgroups in ASD. In ADHD, 5 subgroups have been found that differ in connectivity of fronto-parietal regions (Gates, Molenaar et al. 2014), whereas 3 subgroups emerged that differed in connectivity of reward-related networks (i.e., nucleus accumbens-whole brain connectivity Costa Dias, Iyer et al. 2015). We previously showed that three subgroups exist amongst children with ASD, ADHD, and TD children differing in their levels of cognitive flexibility (Dajani, Llabre et al. 2016), therefore, we predict *at least* three subgroups will emerge that differ on network connectivity profiles important for implementing cognitive flexibility.

Following the results of our previous study (Dajani, Llabre et al. 2016), we hypothesize that one subgroup will include predominantly older TD children with average to above average EFs and low levels of psychopathology. This subgroup may exhibit network connectivity that resembles mature, adult-like networks, characterized by strong, positive connectivity within the FPN, negative functional connectivity between the DMN and FPN (Fair, Nigg et al. 2012, Satterthwaite, Wolf et al. 2013), and integration between SN, FPN and subcortical nodes (Marek, Hwang et al. 2015, Morgan, White et al. 2018). We expect another subgroup to include predominantly younger TD children with less mature network profiles, including weaker within-network connectivity (Bassett, Xia et al. 2018), stronger between-network connectivity, and weaker integration between task-positive networks. In line with the delayed maturation hypothesis (Rubia 2018), we expect this group to also include older children with ADHD who have intact EFs. Children with ASD without elevated comorbid ADHD symptomatology may also comprise this subgroup. Finally, we expect a third subgroup will include children with ADHD and ASD with impaired EFs, elevated general psychopathology (Martel, Pan et al. 2017) and aberrant modular architecture of brain networks (Xia, Ma et al. 2018), which may present as weak connectivity within the FPN and higher FPN-DMN connectivity (Zhong, Rifkin-Graboi et al. 2014, Stevens, Pearlson et al. 2018).

## Methods

### Participants

Participants ages 8 to 13 years (*N*=132, Table 1) included a subset of children used in our previous study investigating heterogeneity in EF ability in TD, ADHD and ASD groups (Dajani, Llabre et al. 2016). Written informed consent was obtained from all legal guardians and written assent was obtained from all children. All procedures were approved by the Institutional Review Board at the Johns Hopkins School of Medicine and all methods were carried out in accordance with the approved guidelines.

**Table 1.** Sample demographics

|  | Diagnostic groups |  |  | *P*<br>value |
| --- | --- | --- | --- | --- |
|  | TD<br>*n*=53 | ADHD<br>*n*=43 | ASD<br>*n*=36 |  |
| **Sex** | 39 M/14 F | 32 M/11 F | 28 M/8 F | .90 |
| **Age** | 10.37 (1.04) | 9.93 (1.22) | 10.52 (1.38) | .09 |
| *range* | [8.00 - 12.58] | [8.00 - 12.33] | [8.00 - 12.92] |  |
| **Race<sup>a</sup>** | 7, 4, 9, 33 | 7, 0, 8, 28 | 2, 0, 3, 29 | .10 |
| **Ethnicity, No.**<br>**Hispanic/Latino** | 1 | 5 | 4 | .14 |
| **FSIQ<sup>b</sup>** | 118.23 (13.25) | 109.00 (11.95) | 103.17 (12.45) | <.001 |
| *range* | [90 - 147] | [87 - 136] | [73 - 131] |  |
| **Motion<sup>c</sup>** | 0.17 (0.09) | 0.22 (0.12) | 0.21 (0.12) | .11 |
| **Handedness<sup>d</sup>, No. L,A,R** | 4, 1, 47 | 5, 0, 38 | 4, 0, 31 | .74 |

a: Numbers for each of the following racial categories presented in the following order: African American, Asian, Biracial, Caucasian, b: FSIQ: WISC-IV full-scale IQ, c: Mean framewise displacement for raw rs-fMRI data calculated in FSL, d: Number of children with left, ambidextrous, right handedness.

### Diagnostic and neuropsychological measures

Community diagnoses of ASD were confirmed with the Autism Diagnostic Observation Schedule (ADOS-G, Lord, Risi et al. 2000) or ADOS-2, (Lord, Rutter et al. 2012), based on study enrollment date) and Autism Diagnostic Interview-Revised (ADI-R, Rutter, Le Couteur et al. 2005). All ASD participants scored ≥7 on the total score on the ADOS-2 or the communication and social interaction score on the ADOS-G. All ASD participants with data for the ADI-R (*n*=34, 94% of ASD sample) met criteria for ASD based on established cutoffs on the ADI-R (≥10 for social interaction, ≥8 for communication/language and ≥3 for RRBs) except for one ASD participant. This participant was still included in the ASD group because they met criteria based on the ADOS-G (communication and social interaction score: 13).

The Diagnostic Interview for Children and Adolescents IV (Reich, Welner et al. 1997) was used to confirm community ADHD diagnoses, determine whether children with ASD had comorbid ADHD, and for exclusionary purposes. Community diagnoses of ADHD were also confirmed with the Conners’ Parent Rating Scales (CPRS-R:L, Conners 1997) or CPRS-3, (Conners 2008), based on study enrollment date) and the ADHD Rating Scale IV, Home version (DuPaul, Power et al. 1998). All participants with ADHD met criteria based on the DICA-IV, except for one child who had missing data. This child met criteria based on both the CPRS-3 and the ADHD Rating Scale IV. TD participants all had T-Scores <65 on both the Hyperactive/Impulsive or Inattentive scales of the Conners’ and only met criteria for ≤3 symptoms on either the Hyperactive/Impulsive or Inattention scales of the ADHD Rating Scale IV. In accordance with the RDoC framework, participants with comorbid psychiatric disorders were not excluded. See Table 2 for detailed diagnostic information.

**Table 2.** Diagnostic information

|  | Primary diagnosis |  |  |
| --- | --- | --- | --- |
|  | TD<br>*n*=53 | ADHD<br>*n*=43 | ASD<br>*n*=36 |
| **Secondary Dx, No. (%)** | **1 (1.9%)<sup>a</sup>** | **20 (46.5%)** | **30 (83.3%)** |
| ADHD-I | 0 | 8 (18.6%) | 9 (25.0%) |
| ADHD-C | 0 | 34 (79.1%) | 12 (33.3%) |
| ODD | 0 | 18 (41.9%) | 9 (25.0%) |
| Simple Phobia | 1 (1.9%) | 5 (11.6%) | 8 (22.2%) |
| GAD | 0 | 0 | 5 (13.9%) |
| OCD | 0 | 0 | 4 (11.1%) |
| Dysthymia | 0 | 0 | 1 (2.8%) |
|  | 0 | 0 | 0 |
| CD,MDD,Mania,Panic,Som,SepAx<sub>b</sub> |  |  |  |
| **ADHD Measures<sup>c</sup>, *M* (*SD*) [*range*]** |  |  |  |
| Conners Hyper/Impulsive | 47.79 (5.51) | 70.55 (12.54) | 66.22 (10.85) |
| Conners Inattention | 45.35 (4.71) | 73.02 (8.2) | 66.78 (10.7) |
| Conners 3 Hyper/Impulsive | 44.93 (7.15) | 75.50 (11.74) | 80.00 (9.55) |
| Conners 3 Inattention | 43.00 (8.82) | 77.20 (9.57) | 83.50 (4.85) |
| ADHD Hyperactivity | 0.22 (0.59) [0-3] | 4.00 (2.89) [0-9] | 3.76 (2.23) [0-8] |
| ADHD Inattention | 0.17 (0.53) [0-3] | 7.00 (1.91) [2-9] | 5.76 (2.75) [0-9] |
| **ASD Measures, *M* (*SD*)** |  |  |  |
| ADI A | -- | -- | 21.03 (5.91) |
| ADI B | -- | -- | 15.71 (4.99) |
| ADI C | -- | -- | 6.21 (2.09) |
| ADOS-2 Social Affect | -- | -- | 7.06 (3.25) |
| ADOS-2 RRB | -- | -- | 4.41 (1.23) |
| ADOS-G CS | -- | -- | 11.9 (2.90) |
| ADOS-G RRB | -- | -- | 3.42 (1.68) |

<sup>a</sup>One TD participant had a past simple phobia of dogs, <sup>b</sup>No child had a diagnosis of conduct disorder, major depressive disorder, mania or hypomania, panic disorder, somatization disorder, or separation anxiety disorder. <sup>c</sup>Reporting *n*=104 for Conners (1997) and *n*=26 with Conners-3 (2008); missing data on Conners *n*=2; Missing data on ADHD Home Rating Scale IV *n*=3; missing data on ADI for *n*=2 ASD participants; *n*=17 ASD participants have ADOS-2 data and *n*=19 ASD participants have ADOS-G data.

#### CBCL

The Child Behavior Checklist (Achenbach 1991) is a parent-report of children’s emotional and behavioral problems. The social problems subscale was used to index social communication and interaction symptoms that are a hallmark of ASD. This subscale is internally consistent (*α* = .82), test-retest reliable (*r* = .90), and validly distinguish between children with and without an ASD diagnosis (Achenbach and Rescorla 2001, Mazefsky, Anderson et al. 2011). T-scores between 67 and 69 represent the borderline clinical range; T-scores≥70 are considered clinically elevated.

#### RBS-R

The Repetitive Behavior Scale-Revised (RBS-R, Bodfish, Symons et al. 2000) is a parent-report of six domains of RRBs: rituals, sameness, self-injurious behavior, stereotypic behavior, compulsive behavior, and restricted interests. The subscales have poor to good interrater reliability (.55-.78) and test-retest reliability (.52-.96). Total RRBs and a subdomain of RRBs, insistence on sameness, have been shown to be correlated with deficits in cognitive flexibility in ASD (Lopez, Lincoln et al. 2005, Miller, Ragozzino et al. 2015), therefore the total score and sameness indices were used in this study. Higher scores indicate greater impairment.

#### Conners’ PRS

The Conners’ Parent Rating Scales-Revised, Long Version (CPRS-R:L, Conners 1997) is a parent report of children’s ADHD symptoms, oppositional defiant disorder and conduct disorder. Here, we used the T-scores from the DSM-IV inattentiveness and hyperactive/impulsive symptom subscales. Higher scores indicate greater impairment.

#### BRIEF

The Behavior Rating Inventory of Executive Function (BRIEF, Gioia, Isquith et al. 2000) is a parent-report of EF impairment of children 5-18 years of age. All T-score subscales will be used to assess EF impairment: inhibition, shift, emotional control, initiate, working memory, plan/organize, organization of materials and monitor. To avoid redundant analyses, we did not analyze the composite scores, which are combinations of the subscales (Behavioral Regulation Index, Metacognition Index, and Global Executive Composite). The subscales are reliable in normative (*r* = .76-.85) and clinical samples (*r* = .72-.84) and can distinguish clinical populations from TD children (Gioia, Isquith et al. 2000). Higher scores indicate greater impairment, with T-scores ≥65 indicating clinical impairment.

### Data acquisition

Children completed a mock scanning session prior to fMRI data collection to acclimatize them to the scanning environment. rs-fMRI data were acquired for participants on a Phillips 3T scanner using an 8-channel head coil (TR=2.5s, flip angle=70°, sensitive encoding acceleration factor=2, 3mm slices, voxel size= 2.7×2.7×3 mm, 156 volumes). The first 10 volumes were immediately discarded to account for magnet stabilization. For the rs-fMRI data acquisition, children were asked to relax with their eyes open and focus on a crosshair while remaining as still as possible. High-resolution T1-weighted scans were also acquired to facilitate registration of the functional image to standard space (TR=8.0ms, TE=3.7ms, 1mm isotropic voxels). Participants were asked to withhold stimulant medication (e.g., Adderall) on the day of MRI scanning, similar to prior neuroimaging studies comparing children with ASD and ADHD (Di Martino, Zuo et al. 2013, Dennis, Jahanshad et al. 2014). Non-stimulant medications were continued as prescribed (e.g., antidepressants, allergy medication). TD children were not taking any psychotropic medications.

### Preprocessing

There were systematic differences in the length of resting-state scans by diagnostic group (*n*=16 ASD children were scanned using a 128-volume protocol, while only *n*=1 TD and *n*=1 ADHD child were scanned using this shorter scan length protocol). To maximize power to estimate connectivity maps, which is dependent on the length of the timecourse, only participants with the longer protocol (156 volumes) were included in this study. Participants with maximum absolute motion in any of the six rigid directions >3mm/degrees were excluded. Preprocessing was conducted using a combination of FSL 5.0.9 (https://fsl.fmrib.ox.ac.uk/fsl/fslwiki) and SPM12 (http://www.fil.ion.ucl.ac.uk/spm/doc/). First, structural images were brain extracted using FSL’s BET tool. Using SPM12 and custom MATLAB scripts, structural images underwent resampling to the EPI image resolution, coregistration to the subject’s mean EPI, and segmentation into grey matter, white matter (WM), and CSF components. These WM and CSF masks were used to compute average WM and CSF timecourses to be used as nuisance regressors at a later preprocessing step. Using FSL’s FEAT, raw fMRI data underwent motion correction, 4D intensity normalization, smoothing with a 6mm kernel, and estimation of linear and non-linear warping parameters to normalize to the MNI152 2mm template. Next, independent component analysis-based automatic removal of motion artifacts (ICA-AROMA) was used to remove motion-related artifacts in native space (Pruim, Mennes et al. 2015). ICA-AROMA works by running an individual-level ICA for each subject, classifying motion-related components as noise, and regressing out motion-related components’ signals from the individual’s 4D time course. The residual time course (with motion-related signal regressed out) was used for subsequent analyses. The denoised data underwent additional nuisance regression (WM, CSF, and linear trends) and band-pass filtering (.01-.10 Hz). Finally, warping parameters generated at an earlier step using FSL were used to normalize the data to the MNI 152 2mm template.

### Region of interest selection

The ideal set of regions of interest (ROIs) for this study would be regions that are consistently activated when engaging cognitive flexibility in children, identified by a meta-analysis of neuroimaging studies that used psychometrically validated cognitive flexibility tasks. Unfortunately, no current meta-analyses of cognitive flexibility tasks exist that are specific to middle childhood. Further, the meta-analyses of cognitive flexibility that do exist for adults include a mix of explicit and inductive tasks that were not psychometrically validated (e.g., Kim, Cilles et al. 2012). Individual neuroimaging studies of cognitive flexibility in children report inconsistent results regarding regions engaged, and are therefore not ideal to guide ROI selection (Dajani and Uddin 2015). Due to the numerous advantages of the Flexible Item Selection Task (FIST), including its established reliability and validity, we opted to use the activation-based results of a study of the FIST in adults to inform ROI selection for this study (Dajani et al., submitted). Further supporting the FIST’s use in guiding ROI selection, the laboratory-based version of the FIST has been validated in children (Dick 2014) and has previously been used in a study of children with ASD (Yerys, Wolff et al. 2012).

Using an fMRI-adapted version of the FIST, we identified cortical and subcortical brain regions that were activated over-and-above basic visual and motor processes associated with the task (Flexibility > Control contrast). Additionally, to define regions within the DMN, which generally deactivate when engaging in cognitive flexibility, we identified regions that were more active during the visual-motor control than for the flexibility trials (Control > Flexibility contrast). To facilitate network analyses and interpretations, the Power (2011) parcellation (Power, Cohen et al. 2011) scheme was used to assign ROIs to networks. Nodes within four large-scale networks integral to cognitive flexibility were included: frontoparietal network (FPN), salience network (SN), DMN and the subcortical network (SUB) (Niendam, Laird et al. 2012, Vatansever, Manktelow et al. 2016). Specific nodes were chosen based on their activation or deactivation in response to the Flexible Item Selection Task (Figure 1, Table 3). Within the FPN, we included the left inferior frontal junction (lIFJ), bilateral frontal eye fields (FEF) and bilateral superior parietal lobule (SPL). Within the SN, we included bilateral anterior insula (AI) and the dorsal anterior cingulate cortex (dACC). Within the DMN, we included the ventromedial prefrontal cortex (vmPFC), posterior cingulate cortex (PCC), and bilateral temporal poles. Finally, within SUB, we included bilateral globus pallidus and thalamus. The SEM-based connectivity analysis utilized here (GIMME) does not perform well with large numbers of ROIs. Accordingly, to reduce the total number of ROIs included, cingulo-opercular, dorsal attention, ventral attention, primary sensory and cerebellar networks were excluded from this analysis. ROIs were centered on the coordinates listed in Table 3 with a 4mm-radius sphere. Timecourses were averaged within each ROI and carried forward for estimation of network connectivity maps using the ‘gimme’ package (version: 0.4) in R (version: 3.3.1) (Gates, Lane et al. 2017, Lane and Gates 2017).

**Table 3.**
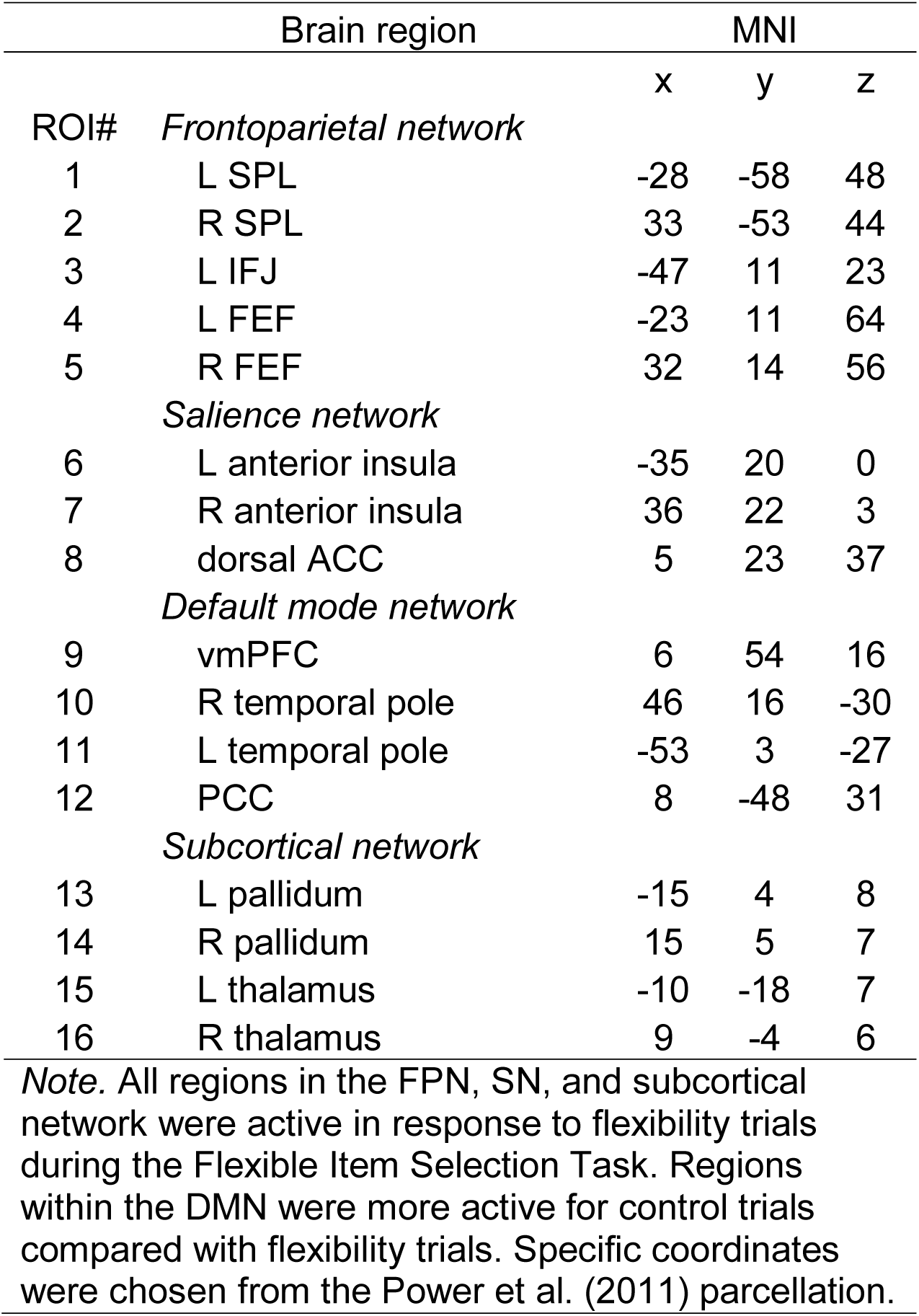
Regions of interest used in GIMME analysis

| Brain region |  | MNI |  |  |
| --- | --- | --- | --- | --- |
| ROI# |  | x | y | z |
|  | *Frontoparietal network* |  |  |  |
| 1 | L SPL | -28 | -58 | 48 |
| 2 | R SPL | 33 | -53 | 44 |
| 3 | L IFJ | -47 | 11 | 23 |
| 4 | L FEF | -23 | 11 | 64 |
| 5 | R FEF | 32 | 14 | 56 |
|  | *Salience network* |  |  |  |
| 6 | L anterior insula | -35 | 20 | 0 |
| 7 | R anterior insula | 36 | 22 | 3 |
| 8 | dorsal ACC | 5 | 23 | 37 |
|  | *Default mode network* |  |  |  |
| 9 | vmPFC | 6 | 54 | 16 |
| 10 | R temporal pole | 46 | 16 | -30 |
| 11 | L temporal pole | -53 | 3 | -27 |
| 12 | PCC | 8 | -48 | 31 |
|  | *Subcortical network* |  |  |  |
| 13 | L pallidum | -15 | 4 | 8 |
| 14 | R pallidum | 15 | 5 | 7 |
| 15 | L thalamus | -10 | -18 | 7 |
| 16 | R thalamus | 9 | -4 | 6 |

*Note.* All regions in the FPN, SN, and subcortical network were active in response to flexibility trials during the Flexible Item Selection Task. Regions within the DMN were more active for control trials compared with flexibility trials. Specific coordinates were chosen from the Power et al. (2011) parcellation.

**Figure 1.**
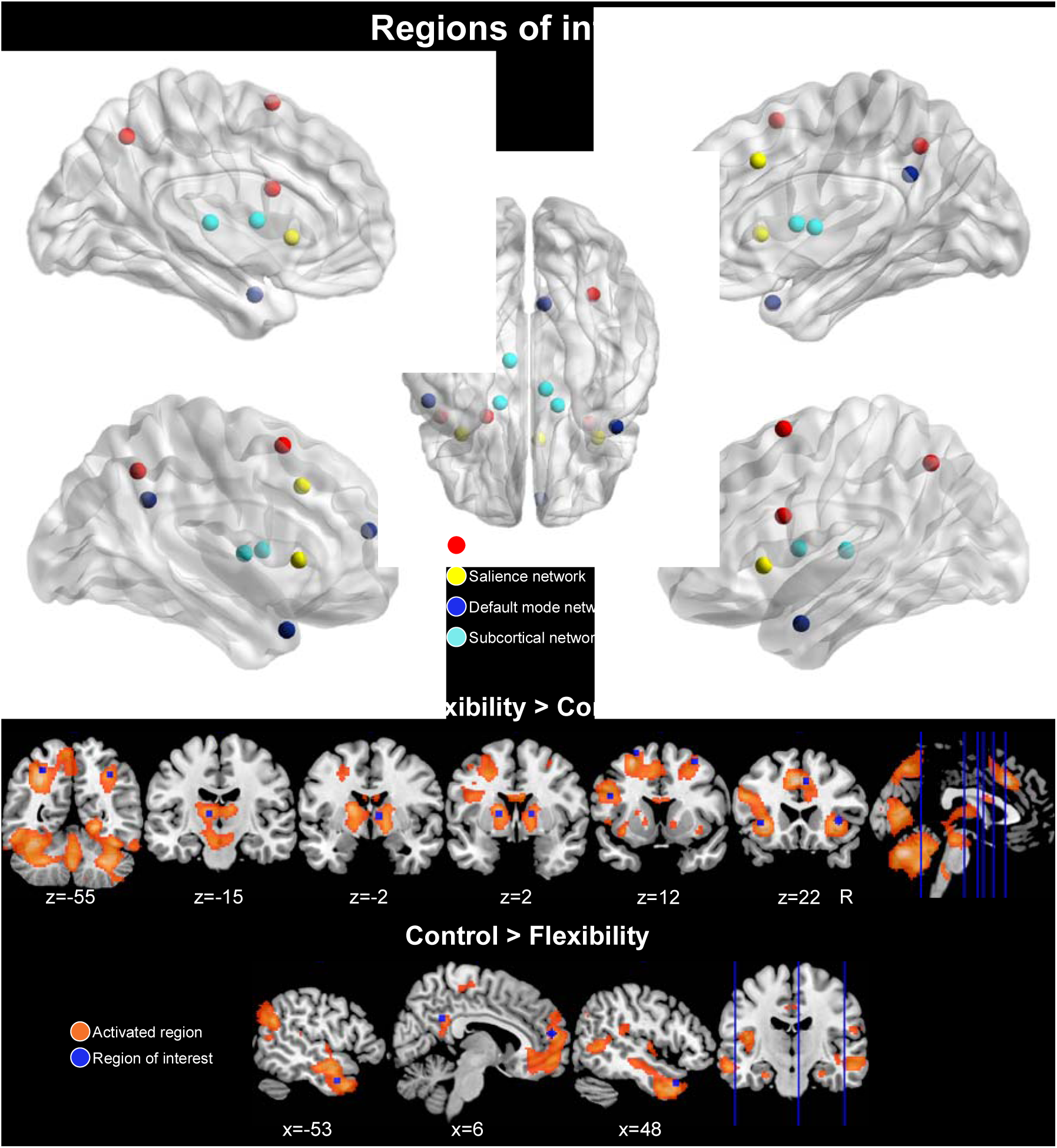
Regions of interest. Sixteen regions of interest used in the GIMME network connectivity analysis. Regions were chosen based on their activation (Flexibility > Control) or deactivation (Control > Flexibility) in response to the Flexible Item Selection Task. Specific coordinates were based on the Power et al. (2011) parcellation to facilitate categorization within large-scale brain networks (frontoparietal, salience, default mode, and subcortical). Brain nodes in upper panel were visualized with BrainNet Viewer (http://www.nitrc.org/projects/bnv/, Cao, Wang et al. 2014).

### Group-, subgroup-, and individual-level connectivity map estimation

GIMME (Gates, Lane et al. 2017) capitalizes on the strengths of the unified SEM framework (uSEM, Kim, Zhu et al. 2007, Gates, Molenaar et al. 2011), which is an extension of SEM to timeseries applications. As part of GIMME, uSEM is used to estimate effective connectivity between pre-specified ROIs on the group-, subgroup- and individual-levels. The uSEM framework incorporates both contemporaneous (*t*) and time-lagged (*t*-1) information between brain regions, which reliably recovers both the existence of a connection and its directionality (Kim, Zhu et al. 2007). Directed connections are estimated contemporaneously, controlling for lagged and autoregressive effects (AR). AR effects indicate the relationship between activity in a single region at time *t* and *t*-1. The benefit of including AR effects, in addition to aiding in reliable estimation of contemporaneous effects for a given brain region, is the ability to estimate path *directionality*. Using a Granger causality framework (Granger 1969), a brain region *η*_1_ is said to Granger-cause activity in another region *η*_2_ if *η*_1_ explains variance in *η*_2_ beyond the variance explained in *η*_2_ by its AR term. Lagged directed connections are also estimated to reduce the chance for spurious contemporaneous effects, but these are not considered to represent underlying neural signal as the temporal resolution of fMRI data is much lower (i.e., seconds) compared to that of neural activity (i.e., milliseconds, Smith, Miller et al. 2011). Instead, when using fMRI data, directional information due to underlying neural signal is best captured in contemporaneous effects (Granger 1969). For this reason, we focus our analysis of network features on contemporaneous effects. The uSEM model is illustrated below, where A is a matrix of contemporaneous effects, Φ is the matrix of lagged effects with AR effects along the diagonal, *η* is the observed time series for a given region of interest, and ζ is the residual for each point in time *t*.

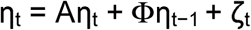

The GIMME algorithm begins by first conducting a group-level search starting with an initial null model, and paths are iteratively added which contribute to better model fit according to multiple modification indices for the majority of participants in the sample, as defined by the user (here, set at 75%). Specifically, GIMME iteratively counts the number of participants whose model would significantly improve if that path were freely estimated, and the path with the highest count is then added to the group-level model. Typically, this process begins by estimating all AR effects first, as this leads to the best performing model search procedure (Gates, Lane et al. 2017). The search procedure continues until there are no paths that would significantly improve the majority of individuals’ models. Next, if the subgroup option is enabled, subgroups consisting of individuals with similar network connectivity patterns are identified using the Walktrap community detection algorithm computed on a sparse count similarity matrix that takes into account the presence of a path and its sign (i.e., positive or negative). The sparse count similarity matrix decidedly outperforms correlation-based similarity matrices according to simulation studies (Gates, Lane et al. 2017). Subgroup-level maps are constructed using an iterative path-adding approach similar to the group-level approach, using the group-level effects as a prior. GIMME adds paths that improve the model for the greatest number of individuals in the subgroup, which must be at least the majority of the sample (here, 51%). Simulation studies demonstrate that given a sample size of at least 75 participants, GIMME accurately recovers up to four subgroups even in cases where subgroup size is unequal (Gates, Lane et al. 2017). The final step estimates individual-level models by adding any additional paths to the group- and subgroup-level paths to best explain that individual’s data. The individual-level model search procedure stops after meeting criteria for excellent model fit for two of four fit indices: comparative fit index (CFI≥.95), non-normed fit index (NNFI≥.95), root-mean-square error of approximation (RMSEA≤.05), and the standardized root-mean-square residual (SRMR≤.05) (Brown 2006). These criteria were also used to identify good individual-level model fit in the current study.

### Cluster validation

To determine the validity of the cluster solution arrived at using the Walktrap hierarchical clustering algorithm within the GIMME framework, stability and validity of the cluster solution was evaluated using the R package perturbR (Gates, Fisher et al. 2018). This algorithm incrementally introduces noise to network edges while maintaining the original graph’s overall properties and compares resulting cluster solutions with the solution for the original network (i.e., using the full sample). A stable solution will not change drastically given small changes to the network. Quantitatively, a cluster solution is said to be stable if the graph had 20% or more of its edges perturbed before the cluster solution for the rewired graph is as different as when 20% of the nodes are randomly placed into different clusters. This is quantified by two distinct, but complementary, metrics that describe the degree to which two community solutions differ: Hubert-Arabie Adjusted Rand Index (ARI) and Variation of Information (VI). To ensure the subgrouping solution is not simply capitalizing on chance where no true subgroups exist, a relative measure of cluster solution quality (i.e., modularity) was used to compare the original cluster solution’s quality with a solution obtained from a random graph that contains no clusters. The cluster solution was considered valid if modularity for the original solution is greater than or equal to the 95^th^ percentile of modularity obtained from random graphs.

### Subgroup characterization: network features

Multiple network metrics were calculated for each individual derived from their connectome data produced by GIMME. Prior work implicates the right anterior insula (rAI) as a hub of causal outflow, interacting with the dACC, dorsolateral prefrontal cortex (dlPFC), ventrolateral prefrontal cortex (vlPFC), and posterior parietal cortex (PPC) (Uddin, Supekar et al. 2011, Supekar and Menon 2012). Therefore, we assessed subgroup differences in out-degree of the rAI normalized by the in-degree (i.e., the number of nodes with which the rAI has an ‘outward’ contemporaneous connection minus the number of contemporaneous connections that terminate on the rAI). Out- and in-degree was calculated using the R package ‘igraph’ (Csardi and Nepusz 2006). To characterize whether subgroups were “hyperconnected” or “hypoconnected” relative to one another, total number of contemporaneous connections were compared. Within- and between-network connectivity was calculated as the number of contemporaneous ROI-ROI connections within- and between-networks, respectively. Integration was indexed with a graph metric called the participation coefficient (PC), which describes the relative distribution of between- and within-network connections for a given node (Guimera and Amaral 2005). A PC approaching 1 indicates that the node’s connections are evenly distributed among all networks and a PC of 0 indicates that all of a node’s connections are within-network. PC was calculated for each node and averaged within network, resulting in a mean PC value for each network, following the methods of a recent network development study (Marek, Hwang et al. 2015). PC was calculated using the R package ‘brainGraph’ (Watson 2018).

## Results

### Preliminary analyses

To quantify the relationship between in-scanner head motion and demographic variables of interest, we characterized the relationship between mean framewise displacement FD (Power, Barnes et al. 2012) for data following ICA-AROMA, age, and diagnostic group. A 2×1 repeated-measures ANOVA demonstrated a significant decrease in mean FD following preprocessing, demonstrating the efficacy of this preprocessing pipeline, *F*(1, 260)=33.00, *p*<.001 (raw: *M(SD)*= 0.20 (0.11), AROMA: *M(SD)*= 0.04 (0.02), Figure 2). There was no difference between ASD, ADHD, and TD groups in mean FD for ICA-AROMA-preprocessed data (*F*(2,129)=0.57, *p*=.57). Similarly, there was no association between age and motion following preprocessing (*r*(124)=.01, *p*=.89). Using the full sample (*N*=132), the GIMME-derived subgroups (which did not align with diagnostic subgroups) differed significantly in mean FD for preprocessed data (*F*(1,130)=16.23, *p*<.001). Therefore, GIMME with subgrouping was rerun using a low motion subsample to reduce potential confounding effects of motion on subgroup formation. This subsample (*n*=99) included participants who were at or below the sample’s 75^th^ percentile of mean FD (≤0.239). This analysis resulted in subgroups who did not differ on motion for preprocessed data (*F*(2,96)=0.14, *p*=.87). Below are the results of subgroup-GIMME with the low motion subsample (*n*=99).

**Figure 2.**
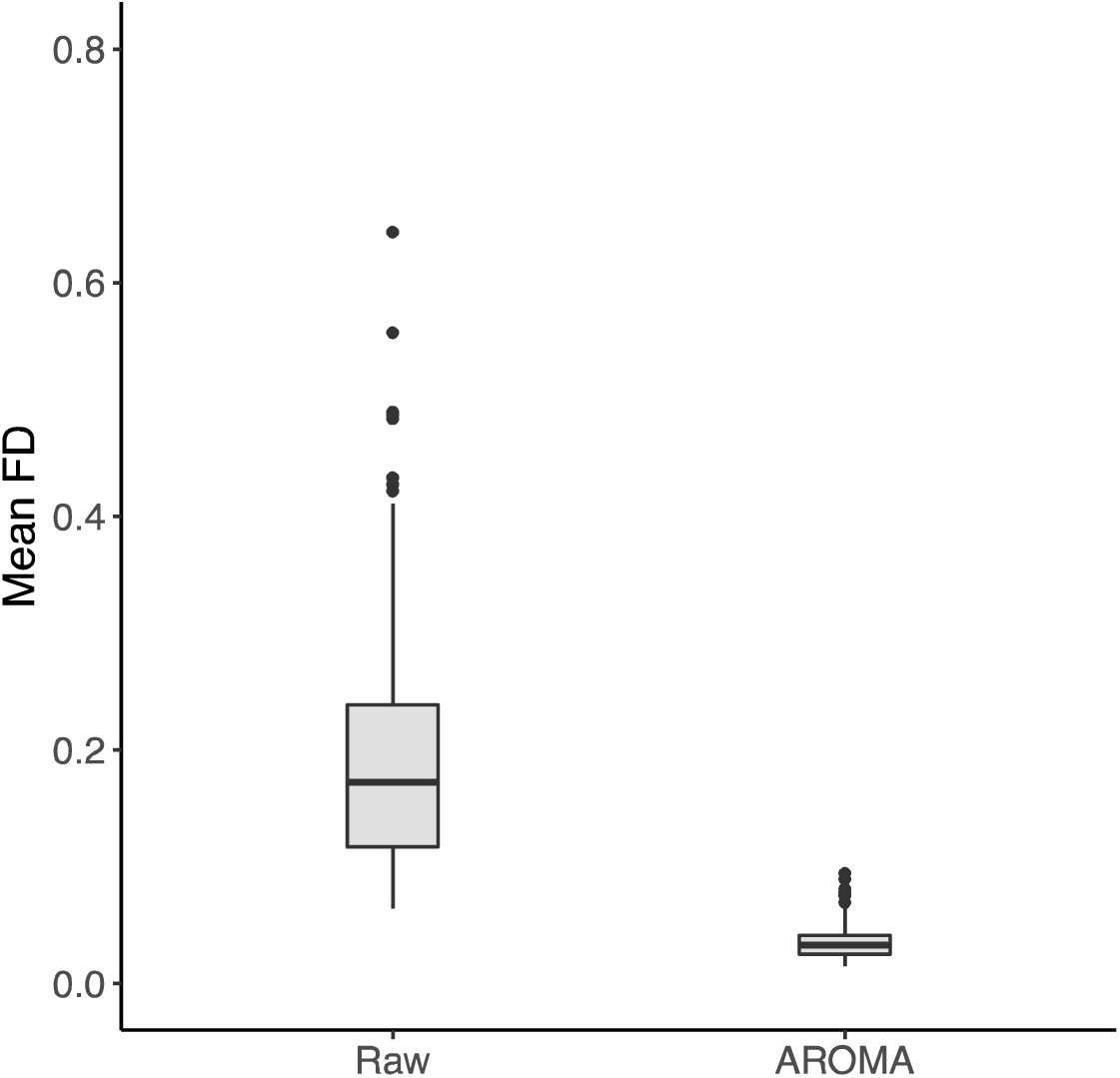
Differences in motion estimates in raw and pre-processed data. AROMA denotes data that has undergone motion artifact removal with ICA-AROMA. Data shown for the entire sample (*N*=132).

### Subgroup-GIMME results using low motion sample

The group-level model included AR effects for all ROIs, one lagged effect from the right pallidum to the right thalamus, and two contemporaneous effects: right pallidum → right thalamus and lSPL → rSPL (Figure 3). These paths were estimated for all 132 participants.

**Figure 3.**
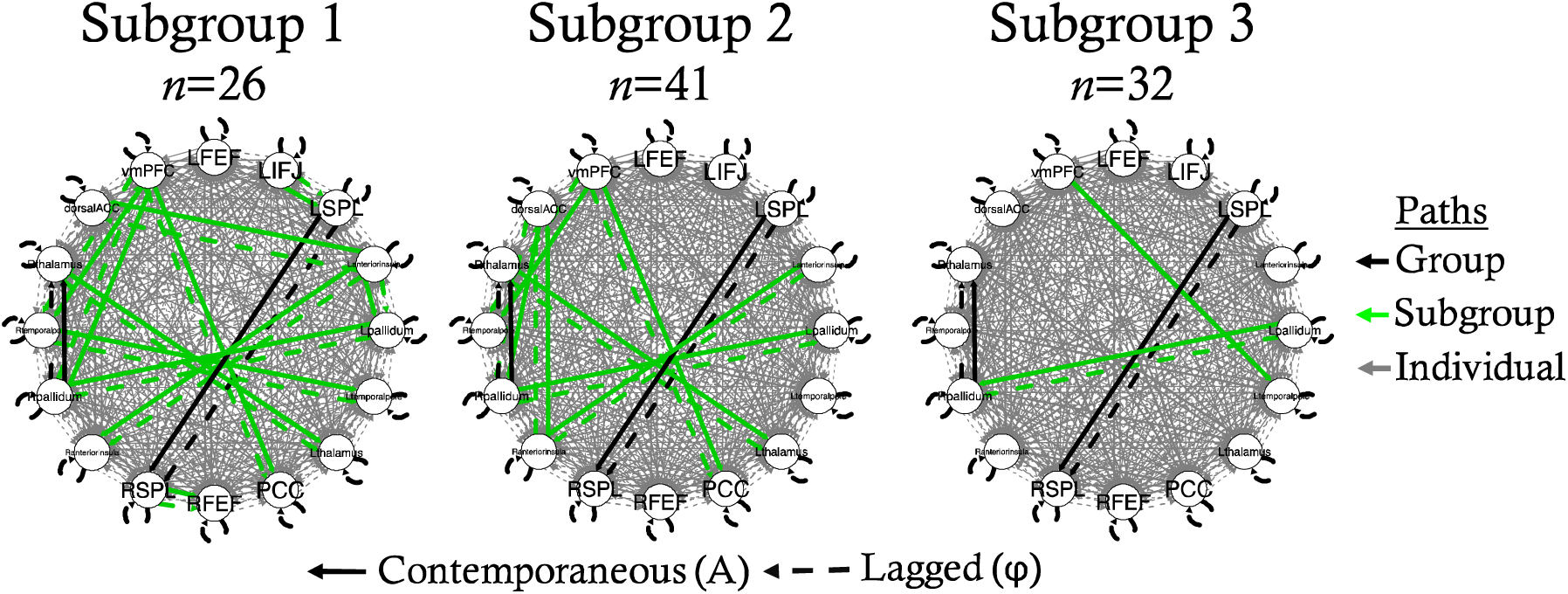
Subgroup-GIMME network models for each subgroup.

#### Model fit

According to approximate fit indices, 98 out of 99 participants’ models had good model fit (CFI: *M(SD)*= 0.95 (0.004), NNFI: *M(SD)*= 0.92 (0.007), RMSEA: *M(SD)*= 0.08 (0.007), SRMR: *M(SD)*=.03 (0.004)). The one remaining participant’s model had excellent model fit according only to SRMR, demonstrating overall acceptable model fit (CFI=.91, NNFI=.86, RMSEA=.11, SRMR=.05).

#### Cluster validation

Using the Walktrap hierarchical clustering algorithm, three subgroups emerged (*n*=26, *n*=41, *n*=32; Figure 3). According to VI, ARI, and modularity, the cluster solution attained was not stable nor valid (Figure 4). Based on VI and ARI, only 1% of edges had to be perturbed before 20% of participants were placed into different clusters than the original solution, demonstrating that minor perturbations to the data caused large changes in the clustering solution (Figure 4a and b). Modularity attained (0.02) was not better than expected by chance (95^th^%ile of random graphs=0.063, Figure 4c), suggesting that clusters are not well defined and participants in different clusters may have more in common than expected if clusters were truly distinct. Based on these results, we concluded that the clustering solution was untrustworthy. Therefore, we do not report subgroup differences in network nor behavioral features.

**Figure 4.**
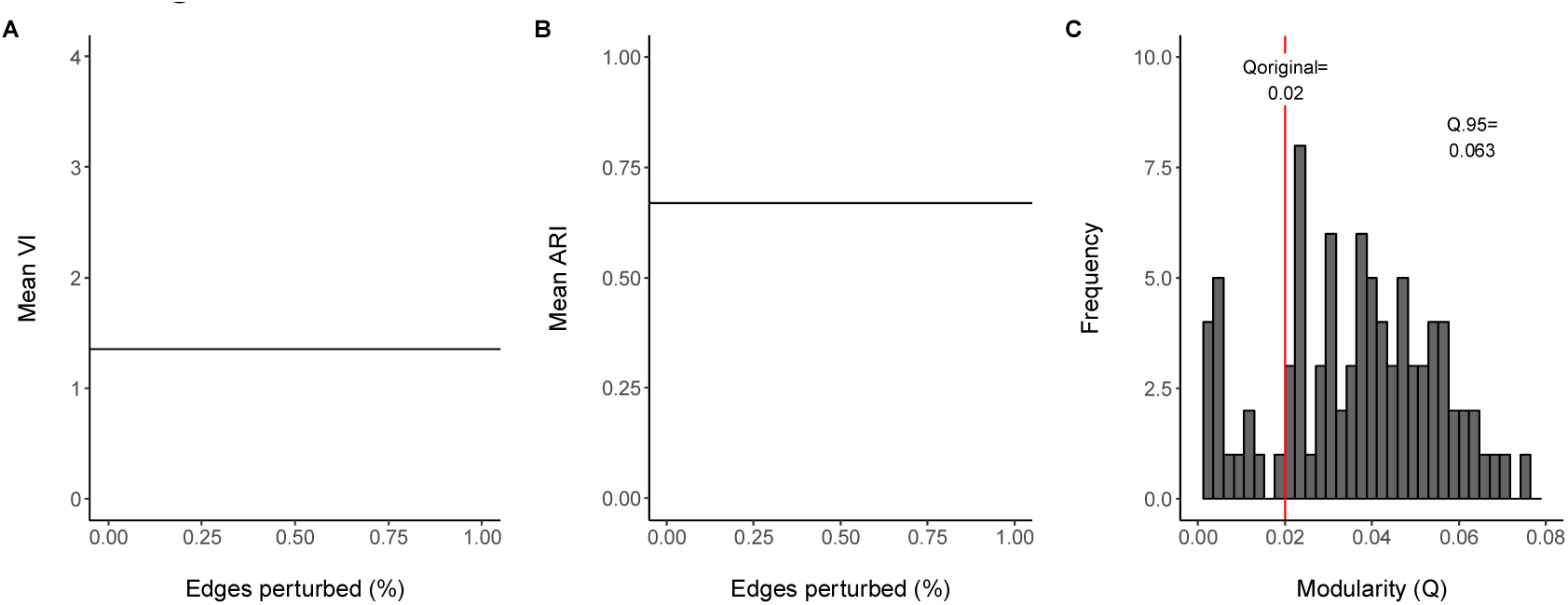
Cluster validation. Results from both VI and ARI demonstrate that the clustering solution is not stable. For panels A and B, the black horizontal line represents the point at which 20% of participants were placed into different clusters than the original solution (20% of nodes perturbed). The dashed vertical line identifies the point at which the perturbed graph reached 20% of nodes perturbed. Black dots represent the perturbed graph based on the original clustering solution while the red dots represent a perturbed random graph. Panel C demonstrates that modularity for the original clustering solution (0.02) was not better than expected by chance (>.06), suggesting this clustering solution is not valid.

## Discussion

To contribute to an improved nosology for neurodevelopmental disorders, we attempted to characterize subgroups of children across ASD, ADHD, and TD groups differentiated by the topography of directed functional connectomes as an alternative to traditional, symptom-based diagnostic systems. Unexpectedly, we were unable to uncover a reliable or valid categorical scheme based on connectomes important for cognitive flexibility in a heterogeneous group of children who ranged from above average to clinical impairment in EF. These unanticipated results highlight the heterogeneity of the topology and strength of connectivity of connectomes important for cognitive flexibility in children with and without neurodevelopmental disorders. Further, these results may suggest that the neural substrates of cognitive flexibility in children may not differ categorically and individual differences in children’s cognitive flexibility may be better represented dimensionally.

The results of the GIMME analysis, which included 16 ROIs and potentially 256 contemporaneous paths to be estimated, identified only two contemporaneous paths common to all 132 participants: from the left to right SPL and from the right pallidum to right thalamus. Unsurprisingly, these two paths are within-network connections (within the FPN and subcortical networks, respectively), which strengthen from middle childhood to early adulthood (Fair, Nigg et al. 2012). Given this result occurred across a heterogeneous sample of children with ASD, ADHD, and TD children, these connections may be foundational to network topology in middle childhood, regardless of psychopathology present. The bilateral SPL group-level path is an example of functional homotopy, which is evident throughout the healthy brain and across the lifespan (Zuo, Kelly et al. 2010). Surprisingly, functional homotopy was not observed for the majority of children in the other bilateral regions included in this study: FEF, AI, temporal poles, pallidum, and thalamus. This suggests that there is heterogeneity in functional homotopy in childhood that may be moderated by levels of psychopathology.

Strikingly, an additional 475 subgroup- and individual-level paths were estimated. The paucity of group-level paths highlights the extreme heterogeneity in network topology among children with and without a diagnosed mental health disorder. This is in contrast with many group-based studies of the development of the *undirected* functional connectome, which conclude that network topology is stable by about 8 years of age (Fair, Nigg et al. 2012, Marek, Hwang et al. 2015). Of note, this result cannot simply be attributed to the mixed diagnostic status of the sample. For example, had all TD children exhibited a similar network topology, a stable subgroup would likely have formed to reflect that. Using a novel individual directed connectome estimation technique, we were able to identify large differences in both within- and between-network topology in middle childhood. These results echo recent calls to regard heterogeneity among healthy and patient populations as not only ubiquitous, but adaptive, due to evolutionary processes which result in many “optimal” brain network profiles (Holmes and Patrick 2018).

The large amount of heterogeneity apparent in the directed functional topography of networks important for cognitive flexibility across a mixed group of children may have impeded the formation of a stable clustering solution in this study, which may be overcome in future studies with a much larger sample size. Simulation studies demonstrate that with a sample size of 75, subgroup recovery is good to excellent using GIMME for up to four subgroups (higher subgroup number was not tested, Gates, Lane et al. 2017). Unfortunately, we had to decrease our sample size from 132 to 99 subjects because the initial clustering solution on the full sample primarily led to clustering based on in-scanner motion. This highlights the importance of considering subgroup differences in nuisance variables such as motion to ensure unsupervised algorithms do not produce subgroups driven by artifacts (Bassett, Xia et al. 2018). Nonetheless, based on the simulation studies it is unlikely that a larger sample size was needed to recover subgroups accurately unless the number of subgroups exceeded 4. This may certainly have been the case, given two DSM-defined diagnostic groups were included, each of which may include multiple subgroups (Georgiades, Szatmari et al. 2013, Stevens, Pearlson et al. 2018). Therefore, the present results do not preclude the existence of 5 subgroups or more within this heterogeneous sample.

Another potential source of the large heterogeneity in functional connectomes observed is spatial variability in the precise location of network nodes across children (Dickie, Ameis et al. 2018). Using templates derived from healthy, young adult samples, Dickie et al. (2018) showed that children show marked variation in the precise location of network nodes and that children with ASD deviate even more than children without a psychiatric diagnosis. Thus, applying ROI coordinates using the Power et al. (2012) parcellation may have led to “missing” true connections due to poor ROI specification on an individual-level, leading to fewer than expected group-level paths.

Despite the above explanations for the unexpected results of this study, a more parsimonious account may be that individual differences in the directed functional connectomes important for cognitive flexibility may best be represented dimensionally instead of categorically. Here, we assumed a categorical structure would best parse heterogeneity in cognitive flexibility in children based on previous studies that identified subgroups present within ASD and ADHD diagnostic categories, which differed in functional connectivity and/or behavioral metrics. For example, subgroups have been shown to exist within children who have ADHD based on differences in functional connectivity of fronto-parietal and reward-related networks (Gates, Molenaar et al. 2014, Costa Dias, Iyer et al.). Subgroups within both ADHD and ASD categories have been demonstrated based on disorder-specific symptoms (Georgiades, Szatmari et al.), neuropsychological task performance (Rommelse, van der Meer et al. 2016, Feczko, Balba et al. 2017), and parent-reports of children’s executive functions (Dajani, Llabre et al. 2016). Moreover, neurodevelopmental disorders are traditionally characterized as distinct, categorical entities, which is practical for clinical translation, where categorical decisions must be made to diagnose and provide treatment (Coghill and Sonuga-Barke).

On the other hand, the traditional categorical approach has recently been challenged by mounting evidence for a dimensional taxonomy of psychopathology (Lahey, Krueger et al. 2017). Studies focusing on parent-report and neuropsychological measures of ADHD symptoms consistently conclude that inattention and hyperactivity/impulsivity are dimensional by nature (Haslam, Williams et al. 2006, Frazier, Youngstrom et al. 2007, Lubke, Hudziak et al. 2009, Marcus and Barry 2011). Further, neuroimaging studies demonstrate that psychopathology, conceptualized as a transdiagnostic ‘p’ factor, dimensionally relates to FPN hypoactivation during a working memory task (Shanmugan, Wolf et al. 2016) and a loss of segregation between the DMN and the FPN and SN at rest (Xia, Ma et al. 2018). Xia et al. (2018) also found a specific relationship between externalizing symptoms (i.e., inattention, hyperactivity/impulsivity, and oppositional defiant symptoms) and stronger SN-FPN coupling at rest. Resting-state fMRI studies focusing on individual disorders have found support for a hybrid categorical/dimensional model in ASD and ADHD based on functional connectivity data (Chabernaud, Mennes et al. 2012, Elton, Alcauter et al. 2014, Elton, Di Martino et al. 2016).

In sum, there is support for both categorical and dimensional models of psychopathology at the behavioral and large-scale neurobiological levels. But, there is a major limitation of the majority of these studies presented, which employ factor or cluster analyses. In these cases, a dimensional or categorical structure is *assumed* to fit the data well, without any formal quantitative tests to determine whether one model is superior to the other. Further complicating matters, clustering algorithms are prone to producing false positives, meaning clusters are produced even in cases where none truly exist. These shortcomings limit the validity of past factor and cluster-based studies of psychopathology. Currently, the only methods that formally determine whether dimensional or categorical models best fit data are taxometric analyses and factor mixture models, which have never before been applied to neuroimaging data, limiting our understanding of how psychopathology should be modeled in consideration of the large-scale neural substrates of behavior. With advances in methodology, future studies may begin to tackle whether a dimensional or categorical model is supported at the brain network level by applying taxometric or factor mixture modeling studies to functional connectome data.

It is important to note that the unreliable and non-modular clustering results presented here are specific to the *a priori* brain regions supplied to the GIMME algorithm. There are many advantages that the GIMME tool has to offer, including individual-level connectome estimation without contamination by group averaging and the estimation of path direction, but this method is not completely data-driven in that only a limited number of ROIs can be used. Thus, researchers are required to use prior literature to guide ROI selection, possibly leading to missed information about functional network topology outside of the networks examined. It is possible that stable subgroups may have been identified using a different set of ROIs, and thus does not preclude the existence of a categorical structure of psychopathology in consideration of other regions and/or functional networks.

Towards an effort to understand whether psychopathology is best represented categorically or dimensionally at the neurobiological level, several future directions are notable. First, in order to construct an alternative nosology that captures the full range of psychopathology, studies should opt to include a wider swath of diagnostic groups to identify relationships beyond ASD and ADHD symptoms, such as frequently comorbid internalizing symptoms (Zald and Lahey 2017). By capturing multiple symptom types, researchers can test whether the neural substrates of psychopathology operate at multiple hierarchical levels from symptoms to the general ‘p’ factor (Zald and Lahey 2017). Considering that neurodevelopmental disorders unfold across age, it may be more fruitful to investigate developmental trajectories in place of cross-sectional studies (Morgan, White et al. 2018). Recent advances in neuroimaging allow for different modalities such as structural, functional, and diffusion-weighted images to be combined in multilayer networks, which may be more informative than one modality alone (Morgan, White et al. 2018).

This study was one of the first to use functional connectome data estimated at the individual level for a heterogeneous group of children spanning TD, ASD, and ADHD diagnoses to test whether the neural substrates of EF and psychopathology follow a categorical or dimensional pattern. Results demonstrated high levels of heterogeneity in the topography of directed functional connectomes important for cognitive flexibility in children with a range of EF abilities. Further, our results did not support a categorical scheme. These results may suggest a dimensional model may better describe individual differences in the neural substrates of cognitive flexibility in children

